# Disturbance of poised chromatin coincides with increased expression of developmental genes in cancer

**DOI:** 10.1101/017251

**Authors:** Stephan H Bernhart, Helene Kretzmer, Frank Jühling, Peter F Stadler, Steve Hoffmann

**Affiliations:** Chair of Bioinformatics, University of Leipzig, Härtelstr. 16–18, D-04107 Leipzig, Germany; Transcriptome Bioinformatics Group, Interdisciplinary Center for Bioinformatics, University Leipzig, Haertelstr. 16–18, D-04107 Leipzig, Germany; Department of Theoretical Chemistry, University of Vienna, Vienna, Austria; LIFE - Leipzig Research Center for Civilization Diseases, University Leipzig, Leipzig, Germany; Max-Planck-Institute for Mathematics in Sciences, Leipzig, Germany; Santa Fe Institute, Santa Fe, New Mexico, USA

## Abstract

Poised (bivalent or paused) chromatin comprises activating and repressing histone modifications at the same location (Voigt et al., 2013). This combination of epigenetic marks keeps genes expressed at low levels but poised for rapid activation (Margaritis and Holstege, 2008; Mikkelsen et al., 2007). Typically, DNA at poised promoters carries low levels of methylation in normal cells (Meissner et al., 2008; Roadmap Epigenomics Consortium et al., 2015), but frequently shows elevated methylation levels in cancer samples (Hinoue et al., 2012; Gal-Yam et al., 2008; Ohm et al., 2007; Rodriguez et al., 2008; Easwaran et al., 2012). Although higher levels of methylation are normally associated with transcriptional silencing, recently counter-intuitive positive correlations between methylation and expression levels have been reported for two cancer types (Hahn et al., 2014; Kretzmer et al.,). Here, we analyze one of the largest combined expression and methylation data-sets to date, comprising over 5,000 samples and demonstrate that the hypermethylation of poised chromatin in conjunction with up-regulation of the corresponding genes is a general phenomenon in cancer. This up-regulation affects developmental genes and transcription factors, including many genes implicated in cancer. From analysis of 7,000 methylation data sets, we built a universal classifier that can identify cancer samples solely from the hypermethylation status of originally poised chromatin. We reason that the alteration of the epigenetic status of poised chromatin is intimately linked to tumorigenesis.

## INTRODUCTION

Poised promoters and enhancers are abundant chromatin states in both stem cells and differentiated cells (Mikkelsen et al., 2007). They are characterized by the simultaneous enrichment of activating (eg. Histone H3 lysine 4 monomethylation [H3K4me1] or H3K4me3) and repressing (eg. H3K27me3) chromatin marks. While the associated genes are repressed, the poised promoters are preloaded with poised polymerase II (Pol II) to prepare genes for rapid activation (Gaertner et al., 2012). Poised chromatin is frequently found in the promoter region of developmentally important genes (Bernstein et al., 2006; Lesch et al., 2013).

In a recent study on the genome wide methylation in malignant lymphoma tissues and normal controls, we showed that the mean methylation change at poised promoters of lymphoma samples is up to three times higher than in other chromatin segments (Kretzmer et al.,). Unexpectedly, the majority of genes controlled by such hypermethylated poised promoters simultaneously show increased expression levels in lymphoma samples. A similar upregulation of genes controlled by poised promoters was reported in colorectal cancers by Hahn et al. (2014). Poised promoters have been suggested to “safeguard differentiation” (Voigt et al., 2013), and their malfunction might have a profound impact on the cell. We therefore investigate whether the *positive* correlation of methylation and gene expression is shared by other cancer types.

## RESULTS

Comparing the 122 non-cancer chromatin segmentations of NIH Roadmap project data (Roadmap Epigenomics Consortium et al., 2015) and the Gencode annotation (Harrow et al., 2012), we confirmed that genes associated with poised chromatin, i.e. poised promoters and poised enhancers, have a highly significant enrichment of the gene ontology (GO)-terms “transcriptional regulator activity” and “developmental process” (see Supplemental Tab. 6-11). While poised enhancers and promoters frequently co-occur, poised promoters are more often associated with protein coding genes (Fisher’s exact test; p<10^−15^) (see Supplemental Fig. 1).

Poising of a gene was not necessarily tissue or cell specific. On the contrary, we identified a large number of frequently poised promoters (FPPs) and enhancers (FPEs) that occur in >50% of all normal cell types. Among the 1,011 FPPs and 2,024 FPEs, the GO-Terms for “developmental processes” and “transcriptional regulator activity” were even more strongly enriched (see Supplemental Tab. 12-17).

Analyzing the relationship between the methylation change of poised chromatin and cancer tissue in the whole genome bisulfite sequencing (WGBS, Supplemental Tab. 1) data of acute lymphoblastic leukemia (ALL) (Lee et al., 2015), we found poised chromatin often to be hypermethylated in ALL. This finding was in stark contrast to the genome-wide hypomethylation observed. In fact, among all types of chromatin segments the strongest increase of mean methylation levels was seen in the poised chromatin (p<10^−16^, Fig. 1A). Strikingly, the contrast between the methylation level of poised segments and the global methylation level was powerful enough to distinguish cancer samples from normal samples. We evaluated 450k methylation arrays of 6,089 cancer and 1,328 normal cells from various tissues (Supplemental Tab. 2-4), and found that the relative methylation of the nine promoter segments poised in most tissues (PPPs, Supplemental Tab. 18-20) separated cancer and normal samples with high sensitivity and specificity (AUC=0.924; Fig. 1B). Despite the high heterogeneity of cancer types (20) and the low number of CpGs (322) in the PPPs, the similarity in the pattern of hypermethylated poised segments served as a robust marker of cancer (see Supplemental Fig. 2,3). A descriptor based on the 12 poised enhancer segments poised in most tissues (Supplemental Tab. 21-23) showed a similar performance, proving that hypermethylation in cancer is a general phenomenon of poised chromatin and not restricted to promoters.

**Figure 1:**
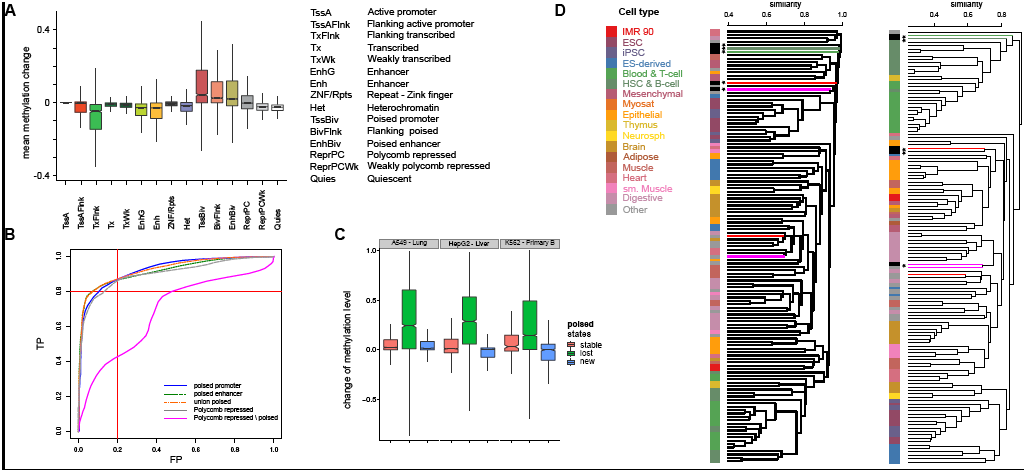
Poised segments in normal and cancer tissues. **A**: Methylation difference for the 15 Roadmap chromatin states in ALL. Poised chromatin shows the strongest hypermethylation. **B**: ROC curve of poised promoter (orange), enhancer (green), union descriptors (blue), polycomb repressed regions (gray) and polycomb repressed regions without frequently poised chromatin (cyan). **C**: Methylation change of poised segments that are stable, are lost or newly occur in cancer cell lines. Data compares cancer cell lines A549 and lung tissue(left), HepG2 and normal liver (center) as well as K562 and Primary B from cord blood (right). **D**: Clustering tree based on the overlap of poised enhancer (left) and active enhancer (right). The tree comprises 127 data sets from Roadmap and ENCODE. In contrast to active enhancers, none of the cancer cell lines (black star) cluster with their cells or tissues of origin when considering enhancers. Colors on leaves correspond to the tissue type, the colors of the edges connect the closest normal tissue to the respective cancer types (magenta: liver, red: lung).

**Figure 2:**
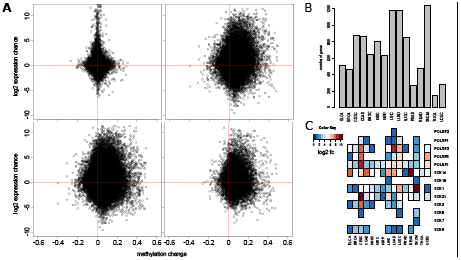
Positive correlation of methylation and upregulation **A:** Scatter plots of methylation change versus expression change for cancer tissue versus normal controls based on the union of 15 cancer data sets. The first quadrant shows up-methylation and higher expression. Top left: data for active promoters (TssA) right: poised enhancers (EnhBiv), bottom left: merged poised intervals right: poised promoters (TssBiv). **B:** Number of positive correlated genes with a fold change of at least 2 for the 15 ICGC cancer types used. **C:** SOX and POU family genes with positive correlation between common poised promoter methylation and expression.

This hypermethylation of poised segments that we observed across all cancer types is difficult to reconcile with the presence of the activating H3K4me3 chromatin mark at these segments (Zhang et al., 2010; Ooi et al., 2007). As H3K4me3 appears to prevent DNA methylation, we hypothesized that the observed hypermethylation is associated with a cancer specific loss of chromatin poising at genome-wide scales. This view was backed by the fact that in three different cancer cell lines, hypermethylation is much stronger in poised segments that were lost than in stably poised chromatin (Fig. 1C).

To further test our hypothesis that cancers have disrupted poised chromatin, we compared the chromatin segmentations of five cancer cell lines from ENCODE (Rosenbloom et al., 2013) and the diverse set of 122 Roadmap normal tissue segmentations. The similarity of two samples was quantified by the overlap of a given type of chromatin segment. As expected, cancer cell lines were similar to their tissue of origin for most chromatin types, including active promoters or enhancers. However, for poised chromatin and Polycomb repressed states, there was no association between the cancer cell lines and their tissue of origin (Fig. 1D, Supplemental Fig. 4,5). The loss of most poised promoters was also described in colorectal tumors (Hahn et al., 2014). Thus, poised chromatin hypermethylation is accompanied by a systematic disruption of poised chromatin in cancer.

To test whether the association of poised promoter hypermethylation with increased gene expression at the corresponding loci is a ubiquitous phenomenon in cancer, we re-evaluated the mean expression values of the RNA-Seq data and 450k arrays of 15 cancer types and their respective controls published by the International Cancer Genome Consortium (ICGC) (International Cancer Genome Consortium, 2010) (see Supplemental Tab. 27)

We found a large degree of positive correlation between increased methylation and expression of associated genes for poised segments. For the union of poised enhancers and promoters, 14 out of 15 cancers showed this positive correlation in more than one third of all cases (Fig. 2A, B, Supplemental Fig. 6,7). 156 of the up-regulated genes showed a fold change (FC) >4 in at least 50% of the cancers. These highly up-regulated genes comprised 78 transcription factors including 53 homeobox genes such as members of the POU (Cook and Sturm, 2008) family as well as SOX (Dong et al., 2004) family genes, which have been linked to tumorigenesis (Abate-Shen, 2002; Haria and Naora, 2013) (Fig. 2C, Supplemental Tab. 28-36).

In accordance with previously published data (Easwaran et al., 2012), cancer cell lines showed a very strong hypermethylation of the poised chromatin states accompanied by the silencing of associated genes (data not shown). On the other hand, data from fresh tissues obtained from the ICGC data base revealed the opposite.

## DISCUSSION

In summary, the rather unexpected and recurrent finding of up-regulated expression co-occurring with hypermethylation of the poised chromatin suggests that (i) the epigenetic poising of RNA polymerase II is lost at some point during tumorigenesis and (ii) the associated genes are expressed. The DNA hypermethylation (iii) may be explained by the enrichment of two DNA methyltransferases (DNMT1 and DNMT3B) surrounding poised promoters in cancer cell lines (Jin et al., 2012). It is tempting to speculate that poised chromatin methylation is a backup to re-enforce repression in case of a disruption of poising, emphasizing the importance of keeping the respective genes under tight control. This need for tight control might also explain the fact that poised chromatin tends to be more enriched in conserved elements than its respective active chromatin counterpart (Roadmap Epigenomics Consortium et al., 2015). In normal cells the stalled RNA polymerase II at poised segments also prevents most transcription factors from binding (Roadmap Epigenomics Consortium et al., 2015). A change of chromatin state is also essential for hypermethylation, since the presence of H3K4me3 would still protect the DNA from methylation.

A simple sequence of events (Fig. 3) can explain the coincidence of these three findings. Initially, the loss of the poised state of chromatin allows for transient activation of a large number of genes. Now, the formerly poised intervals are accessible for chromatin remodeling and methylation. Subsequently, chromatin remodeling or DNA methylation re-silences the genes. The re-silencing does not happen instantaneously on a genome wide scale. Accordingly, the transient expression is detectable in heterogenous sets of cancer tissues, but can no longer be found in homogeneous cancer cell lines.

**Figure 3:**
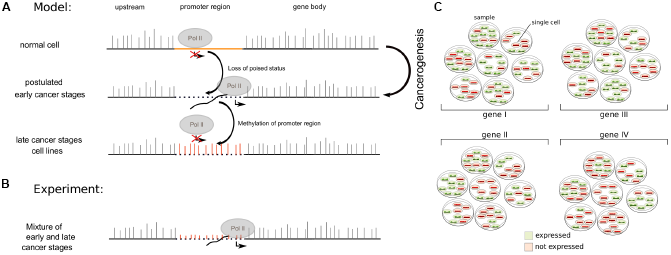
Model for tumorigenesis. **A:** (top) Situation in normal cell: Poised regions show low methylation but no expression due to the stalled polymerase. (center) Postulated early cancer stage: Loss of poising leads to expression, methylation is not yet sufficient to repress expression. (bottom) Late cancer stage: Methylation or chromatin remodeling has repressed expression from the formerly poised gene, e.g. seen in cancer cell lines. **B**: Situation seen in experiments: Mixture of patient samples and cells within samples show partial expression and methylation. **C**: Illustration of situation in fresh cancer tissues. Every patient (Petri-dish) shows a heterogeneous set of expression of previously poised genes. In some of the cancer cells, they are still expressed (green), while in others repression has already happened (red). The expression status is different for every gene, leading to the situation seen in the experiments in b.

One probable trigger of disruption of poising could be the proto-ongogene MYC. MYC is frequently over-expressed in cancer (Beroukhim et al., 2010) and has been predicted to cause expression of formerly poised genes (Rahl et al., 2010). However, the exact mechanism of MYC-induced tumorigenesis is not yet known. We propose, that by initiating the transcription of many poised developmental genes, MYC might contribute to the de-differentiation of cells and consecutively to cancer development. Although we have shown that these alterations are not limited to cancers over-expressing MYC (see Supplemental Tab. 5).

Poised chromatin is known to play an important role in normal tissue development, but also seems to be crucial for keeping cells in their differentiated state. We showed that tumorigenesis in various cancer types is accompanied by the aberrant expression of developmental genes no longer controlled by poised chromatin. Developmental genes play important roles in cell differentiation. Accordingly, the runaway expression of many developmental genes might well lead to de-differentiation of cells, much like the over-expression of the four Yamanaka factors used to generate induced pluripotent stem cells (Takahashi and Yamanaka, 2006). If apoptosis or proliferation related genes are damaged, such de-differentiated cells might develop into cancer cells.

Elucidating the molecular mechanisms behind this loss of the poised state has the potential to reveal novel preventive approaches against cancer. In the short term, measuring the methylation of a small set of loci corresponding to frequently poised chromatin provides a cost-effective and efficient universal cancer descriptor for diagnostic purposes. Identification of prevalently poised segments for special tissues can result in descriptors for sensitive and selective clinical diagnosis in cancers where other molecular descriptors are still lacking.

## METHODS

### Data Acquisition

We downloaded 127 15-state chromatin segmentation annotations from the NIH Roadmap Epigenomics Project (16 of which generated from ENCODE data).

We used normal and cancer RNAseq data from ICGC as well as 450k methylation data from ICGC (International Cancer Genome Consortium, 2010), ENCODE (Rosenbloom et al., 2013) and GEO (Barrett et al., 2013). In total, we retrieved 7,870 cancer 450k data sets and 1,691 normal cell 450k data sets. For the 15 ICGC data sets where we found RNAseq data for normal tissue, we found a total of 390 normal and 5,389 cancer data sets.

### Poised intervals

Poised intervals and their subsets are based on the 122 normal cell chromatin segmentations from Roadmap. Bedtools (Quinlan and Hall, 2010) was used to merge the poised promoter and enhancer segments as well as their union to generate the poised intervals. We counted how many distinct chromatin segmentations contributed to each interval. We aimed to construct small prevalently poised sets of roughly the same size. Thus, the nine poised promoter intervals containing contributions from more than 97 (80%) segmentations (PPP), the 12 poised enhancer intervals containing contributions from more than 115 (95%) segmentations (PPE) and the 14 intervals containing contributions from more than 119 segmentations (98%) for their union (PPM) made up our prevalently poised data sets. For comparison, we also generated a prevalent set for the polycomb repressed (ReprPC) chromatin segments, constructed from the 10 intervals containing data from more than 99% of the normal Roadmap data sets. Furthermore, we generated frequently poised intervals that contained poised segments from at least 50% of the data sets. This resulted in 1,011 promoter (FPP), 2,024 enhancer (FPE) and 3,805 frequently poised intervals from the union of poised enhancer and poised promoter data (FPM). The intervals generated were intersected with the Gencode v19 gene annotation while each gene was extended by 1,500 nt upstream. The resulting genes were investigated for enrichment in GO-terms, KEGG pathways, and other descriptors using GOrilla and DAVID.

### Clustering of chromatin segments

We used bedtools (Quinlan and Hall, 2010) to intersect the different Roadmap annotations. The fraction of segments overlapping with segments of the same type was used to compute correlation matrices using R (R Core Team, 2013). The same fraction data was translated into a distance and used to cluster the datasets using R (using hclust, mcquitty and the mean distances). Mean overall distances between the single segmentations of the tissues were computed using R, and mean distances of cancer to normal cells and normal to normal cells were compared using R’s Wilcoxon function.

### Methylation change of chromatin segments in ALL

We downloaded the ALL data set containing methylation differences from GEO (GSE56601), and took the mean of the “ETV6-ALL difference from preB” and the “HD-ALL difference from preB” values as the methylation difference per CpG. We intersected it with Roadmap’s “Primary B cells from cord blood” segmentation using bedtools and computed the mean methylation difference per interval using bedtools merge. These methylation differences were plotted and analyzed using R.

### Methylation change of poised segments in ENCODE cancer cell line data

The Roadmap chromatin segmentations of cancer cell lines and their tissues of origin (A549:Lung, K562:Primary B from cord blood, HepG2:Liver) were intersected. All segments where either cancer or tissue of origin had poised chromatin were kept. The segments were binned into “stably poised”, where both normal and cancer tissue show poised chromatin (poised enhancer, poised promoter or flanking poised), “poised lost”, where only the normal chromatin was poised, and “poised gained”, where only cancer tissue showed poised chromatin. For methylation analyses, WGBS (lung, liver) and reduced representation bisulfite sequencing (RRBS) data (A549, K562, Primary B from cord blood, HepG2) was downloaded from Roadmap. Methylation changes were computed using CpGs that had valid entries in both normal and cancer data sets only. The methylation differences were analyzed using BigWigAverageOverBed of the UCSC tools (Kent et al., 2010) and plotted using R.

### Methylation descriptor on 450k Arrays

We used the methylation as measured by 450k arrays at the 9 PPP segments (containing 322 CpGs represented within 450k arrays), the 12 PPE segments (706 CpGs) and the 14 PPM segments (1,163) as a descriptor telling normal from cancer samples. Methylation rates were computed using UCSC tools for bigwig files or in-house tools for GEO and ENCODE array data. The identifiers of the array’s CpGs were mapped to the prevalently poised intervals, and mean rates were computed when at least 3 CpGs were present. Datasets where, due to lack of CpG information, a mean methylation rate could only be computed for less than 4 intervals were discarded. About 6,090 cancer and 1,300 normal data sets met these criteria. We used the mean of the mean β-values of the intervals to compute the mean interval methylation rates. We normalized using the mean methylation over all CpGs. For comparison, we used the merged polycomb repressed (ReprPC) data-set (containing 1,649 CpGs in 450k) as well as a ReprPC data set depleted of possibly poised segments. To generate this, we used bedtools to subtract the PPM from the ReprPC data-set. The resulting segments contained 60% of the total nucleotides of the merged ReprPC segments, but only 212 (12.8%) CpGs contained in 450k arrays. Receiver operator characteristics (ROC) curves were plotted using R, area under these curves (AUCs) were computed using R’s AUC package (Ballings and den Poel, 2013).

### Methylation and Expression analysis of cancer data

For correlation of expression and methylation, we used our frequently poised segments (FPP, FPE and FPM) and the genes associated with them. The methylation of all poised segments that could be connected to a gene was computed using UCSC tools. Only poised promoters with at least 3 covered CpGs were kept. Expression of the genes was taken from the RNAseq data downloaded from ICGC. Mean expression was computed over all normal and cancer data sets of the resp. cancer type. The log fold change of the mean reads per kilobase per million (RPKMs) was used for the scatter plots. For the figure 2, we used a union of the data from all 15 cancer types. Where available, we also performed the analysis with chromatin segmentations from the tissues of origin of the respective cancers (see Supplemental Fig. 7). Positively correlating genes with an at least 4-fold higher expression in cancer tissue were analyzed using GOrilla and DAVID. Genes that showed clear positive methylation change (>0.02) and an at least 2-fold increase in expression where counted. For investigating SOX and POU family genes, the methylation difference of all poised promoter intervals shared by at least 50% of the annotations was correlated to the expression change of the respective genes. Only methylation changes >0.04 and expression fold changes >2 were used for further analysis.

## DATA ACCESS

A full list of accession numbers and URLs for the data sets used in this study is provided in Supplementary Tab. 1-4

## ACKNOWLEDGEMENTS

We thank Bernhard Radlwimmer for helpful discussion. This research was supported by the German BMBF (PTJ grant HNPCCSys 031 6065A), and through ICGC MMML-Seq (01KU1002J), and LIFE (Leipzig Research Center for Civilization Diseases), Leipzig University. LIFE is funded by the European Union, by the European Regional Development Fund (ERDF), the European Social Fund (ESF) and by the Free State of Saxony within the excellence initiative.

### DISCLOSURE DECLARATION

The authors declare no conflict of interest.

### AUTHORS CONTRIBUTIONS

SHB, PFS and SH designed the study. SHB, FJ and HK performed the analyses. SHB, PFS and SH wrote the manuscript. All authors read and approved the manuscript.

